# In-depth 15 H7N9 Human Serum Proteomics Profiling Study

**DOI:** 10.1101/2020.01.06.896829

**Authors:** ZiFeng Yang, Wenda Guan, Shiyi Zhou, Liping Chen, Chris K.P. Mok, Jicheng Huang, Shiguan Wu, Hongxia Zhou, Yong Liu, Malik Peiris, Xiaoqing Liu, Yimin Li, Nanshan Zhong

**Affiliations:** State Key Laboratory of Respiratory Disease (Guangzhou Medical University), National Clinical Research Center for Respiratory Disease, First Affiliated Hospital of Guangzhou Medical University, Guangzhou, 510120, PR China; State Key Laboratory of Respiratory Disease-Research Laboratory for Clinical Virology, National Clinical Research Center of Respiratory Disease-Research Center for Clinical Virology, Kingmed Virology Diagnostic & Translational Center, Guangzhou Kingmed Center for Clinical Laboratory Co., Ltd.; Technology Centre, Guangzhou customs district, PR China; HKU-Pasteur Research Pole, School of Public Health, HKU Li Ka Shing Faculty of Medicine, The University of Hong Kong, Hong Kong Special Administrative Region, PR China

**Keywords:** H7N9, avian influenza, cell death, HSPA8, proteomic

## Abstract

**Background:** Human infection by avian influenza viruses is characterized by rapid development of acute respiratory distress and severe pneumonia. However, the underlying host response leading to this severe outcome is not well studied.

**Methods:** We conducted mass spectrometry-based serum proteome profiling on 10 healthy controls and 15 H7N9 infected cases with two time points and carried out statistical and biology functional enrichment analysis.

**Results:** In total, we identified 647 proteins, 273 proteins were only found in H7N9 infected cases which might generate from cell leakage/death (apoptosis and/or necrosis) and identified 50 proteins with statistically significant difference between healthy control and H7N9 infected cases from 168 qualified proteins. We also found that M1 and PB2 tightly associated with the host’s HSPA8 (P11142, *p*=0.0042) which plays an important role in the protein quality control system.

**Conclusions:** H7N9 infection may increase cell programmed/unprogrammed cell death, and we suggested that upregulated extracellular HSPA8 may suppress the H7N9 virion replication via activation amyloid-beta binding network.

## Introduction

As of August 2019, there have been 1568 confirmed human cases and 616 deaths caused by infection with avian H7N9 influenza A viruses in China, with a 39% mortality rate [1]. In April 2019 a fatal case of highly pathogenic avian influenza (HPAI) H7N9 was reported in Inner Mongolia, China, which was associated with severe pneumonia and respiratory failure [2]. Acute respiratory distress syndrome, as observed in the recent human case of HPAI H7N9 infection, may be caused by multiple factors, including higher viral load [3], altered viral tropism and spread in the lower/upper respiratory tract [4], virally-induced host shutoff [5] or hijacking host function [6–9]. Mass spectrometry-based serum proteome profiling has made it possible to conduct extracellular fluid proteins analysis and to investigate important mediators of intercellular communication and host-viral interactions associated with disease progression, coagulation, communication, inflammation and the immune response.

Several studies have been published using mass spectrometry-based proteomics with *in-vitro* assays (Supplementary Table 1)[10–12]. Simon et al. (2015) performed a comparative proteomic study in A549 human cells infected with seasonal H1N1 (sH1N1), pdmH1N1, H7N9 and HPAI H5N1 strains. Using multiplex iTRAQ labelled quantification, they found that NRF2 was associated with infection with HPAI strains of influenza[10]. Ding et al.(2016) carried out similar research work in A549 infected with pdmH1N1 and H7N9 by labelling digested proteins with Cy-dye followed by identification using MALDI-TOF-MS/MS, and found that down-regulation of CAPZA1, OAT, PCBP1, EIF5A were related to the death of cells infected with H7N9, and down-regulation of PAFAH1B2 was related to the later clinical symptoms in patients infected with H7N9[11]. Sadewasser et al.(2017) performed SILAC labelled proteomic assays on human HEK 293T and A549, characterized sets of cellular factors alternatively regulated in cells infected with H3 or H7 AIVs, and suggested that VPRBP(*DCAF1*) was identified as a novel drug target [12]. There has also been a study conducted in humans by Wang et al. (2018), who revealed the differential plasma proteome profiles in two cases of H7N9 infection in a single family[13]. Apolipoproteins appeared to play a critical role in the progression of H7N9 infection and disease. To our knowledge few population-based extracellular fluid proteomic studies on host-viral interactions underlying subtype H7N9 influenza virus infection haves been published.

To better understand host’s response to H7N9 infection, we carried out serum proteomics profiling on 15 confirmed H7N9 human cases at two time points and compared these with 10 healthy controls. These experiments were performed on LC/MS/MS using labelling by iTRAQ method. We also built a local cloud-based analysis workflow for stable isotopic labelled proteomics study to qualify and quantify proteins and conduct comparative analysis. Our study identified key host responses that may be related to the progression of H7N9 disease.

## Materials and Methods

Fifteen patients with laboratory-confirmed, 14 patients in low pathogenic avian influenza A H7N9 infected (LPAI) and one in high pathogenic avian influenza A H7N9 infected (HPAI), were enrolled at First Affiliated Hospital of Guangzhou Medical University, from 2013 to 2017. The clinical history were recorded. The detailed enrollment process has been shown in Figure 1.

**Figure 1:**
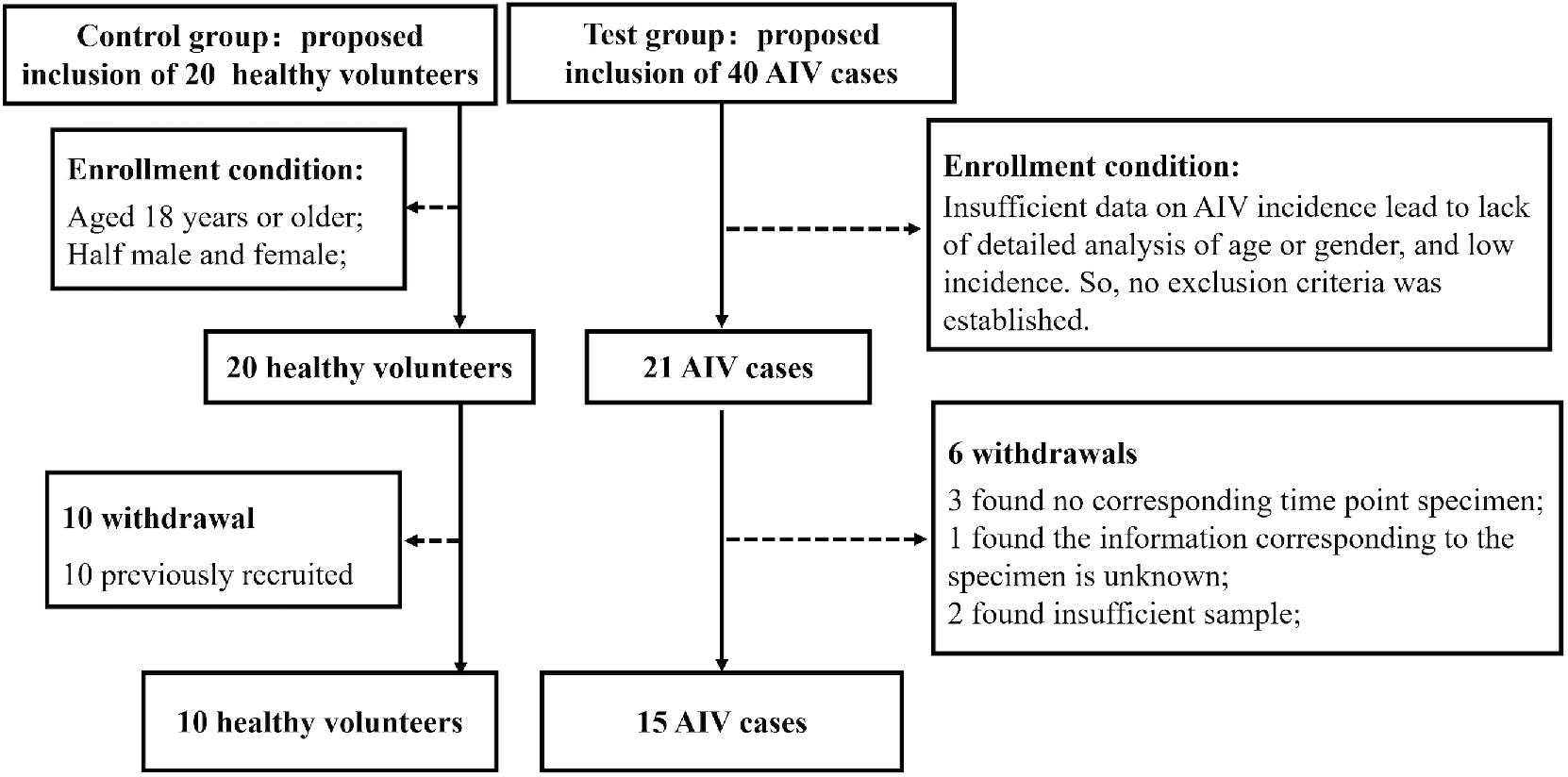
We totally assessed 20 healthy volunteers and 21 H7N9 infected patients, 10 healthy control and 6 H7N9 infected patients withdrawn due to the protocol violations. Finally, 15 patients and 10 healthy volunteers were enrolled in this study.

Earlier time point was defined as the earlier one of the two time points, and Later time point was the later one. The metadata of the patients were tabulated in Supplementary Table 2; and clinical and immunological features of some patients have been reported[3,14]. Ten healthy serum controls were collected from our lab colleagues. Approval for the study was obtained from the ethics committee of the First Affiliated Hospital of Guangzhou Medical University (No.:2016-78) and informed consent was obtained from the patients or their family members.

Proteome profiling of host serum was performed by iTRAQ-based quantitative differential LC/MS/MS, protein identification and quantitation were carried out by in-house workflow deployed in Galaxy framework[15], statistical related analysis was done using MetaboAnalystR (v2.0)[16] in R (v3.6.1), and biological and functional analysis were analyzed by STRING tools[17]. Detailed workflow was provided as supplementary text 1 and showed in Figure 2, we also provided Galaxy workflow for this study as Supplementary workflow 1.

**Figure 2.**
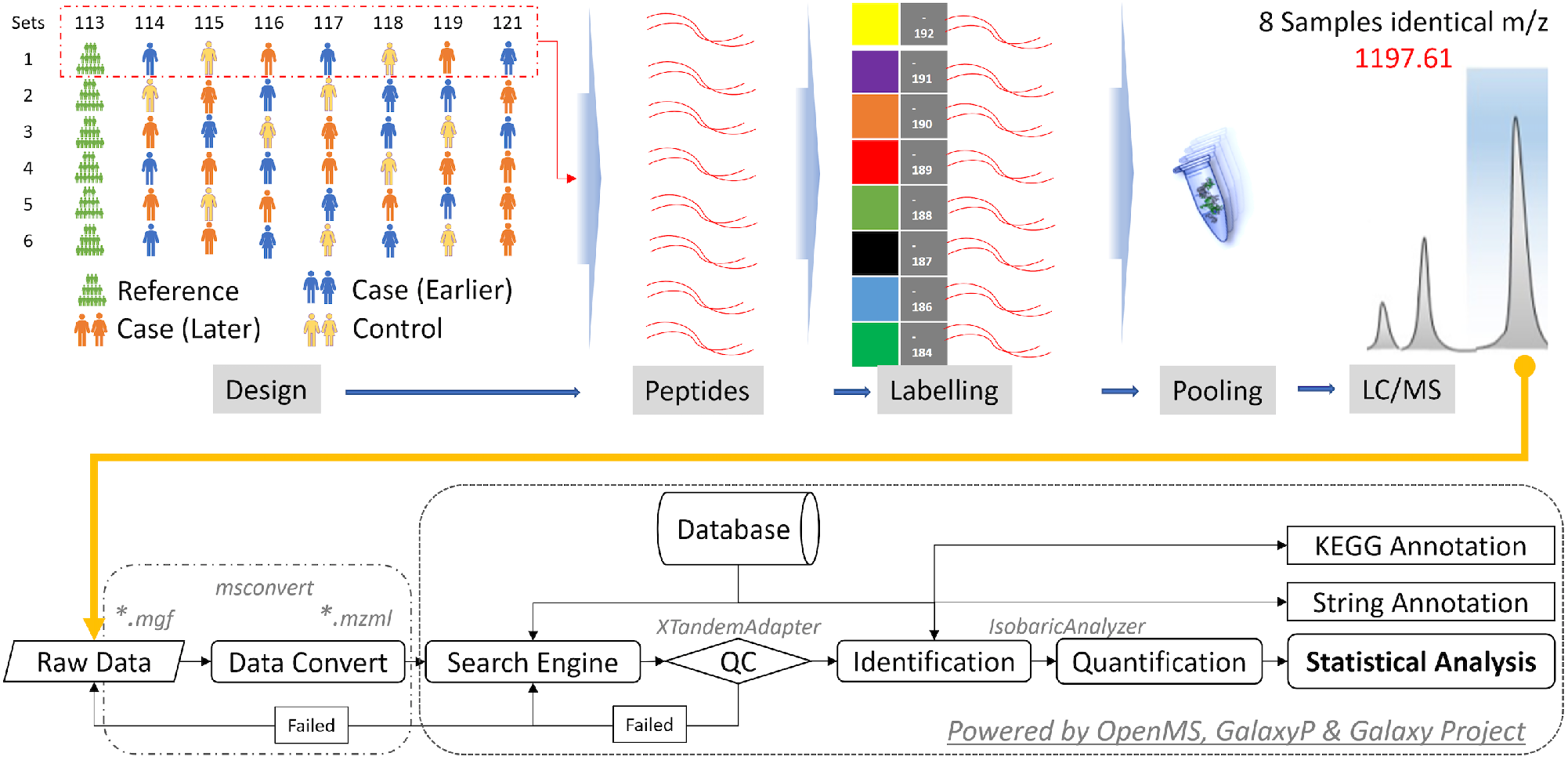
Experimental and analytical scheme of deep depleted serum proteome analysis using iTRAQ LC/MS. H7N9 infected cases and control samples were extracted, depleted high-abundance protein, digested, labeled and pooled, coupled with high-resolution MALDI-TOF/TOF MS (Bruker Daltonics, Billerica. MA). All data were analyzed by the in-house workflow, which was powered by OpenMS, GalaxyP and Galaxy Project.

## Results

### Clinical Characteristics Of the Samples

Fifteen H7N9 infected patients were enrolled in this study from 2013 to 2017. Clinical, virologic, and immunological features of each patient were recorded in detail (Supplementary Table 2), including 14 LPAI and one HPAI patients. Five patients (No. 3, 9, 11, 12, and 14) had a fatal outcome, and the remaining patients were discharged. All samples had laboratory confirmed with H7N9 infection by RT-PCR using specific HA primers. The mean age was (28.60 ± 2.79y) in the control group compared to the case group (49.50 ± 17.83y) with statistical significantly difference. The viral load was determined from the throat swab (termed as an upper respiratory tract specimen) and from BALF or endotracheal aspirate (termed as lower respiratory tract). The mean URT and LRT viral loads level was 2.40 ± 2.14 (Log10), 4.35 ± 1.62 (Log10) in case’s earlier compared to the viral loads 0, 1.04 ± 1.94 (Log10) in later time point with statistical significantly decreased (*p* < 0.05), respectively. We assumed the viral loads in the upper and lower respiratory tracts were zero and the PF ratio (PaO_2_/FiO_2_) values were 450. The mean PF ratio was significantly lower in H7N9 infected cases, and the PF mean value was significantly upregulated after treatment. Table 1 was listed the characteristics of the 25 participants enrolled in this study, including the 10 healthy controls.

**Table 1:** Clinical Characteristic of the groups

|  | Control | Case (Earlier) | Case (Later) |
| --- | --- | --- | --- |
| Size (n) | 10 | 15 |  |
| Gender<sup>*</sup> (Male) | 40% | 53.3% |  |
| Age (Year)<sup>†</sup> | 28.60 ± 2.79 | 49.50 ± 17.83 |  |
| URT (Log10)<sup>#</sup> | NT | 2.40 ± 2.14 | 0 |
| LRT (Log10)<sup>#</sup> | NT | 4.35 ± 1.62 | 1.04 ± 1.94 |
| TWC (x10<sup>9</sup> cells/L) | NT | 11.75 ± 6.55 | 12.33 ± 8.16 |
| CRP (mg/L) | NT | 86.00 ± 44.12 | 70.69 ± 56.81 |
| Procalcitonin (ng/ml) | NT | 1.87 ± 3.68 | 5.49 ± 14.52 |
| PF <sup>#</sup> | NT | 146.79 ± 89.23 | 228.49 ± 168.07 |

<sup>\*</sup>: Male of total group size (%); Value: mean ± SD; NT: Not Tested; URT: Viral Load on Upper Respiratory Tract; LRT: Viral Load on Lower Respiratory Tract; TWC: Total White Cells; CRP:C-Reactive Protein; PF: PO<sub>2</sub>/FiO<sub>2</sub> ratio
<sup>†</sup>: Significantly difference between case and control group with *p* value less than 0.05 <sup>#</sup>: Significantly difference between earlier and later time point in case with *p* value less than 0.05

### Overview of H7N9 Infected and Healthy Control Sera Proteomics Profiling

Based on 130,325 qualified and quantified MS/MS spectra (FDR < 0.01) in all samples, 36,708 non-redundant peptide hits were identified. Identification rates were approximately 28.17% and a total of 647 proteins had been annotated. The detailed information regarding the peptide and protein identification and quantification of six sets generated by X! Tandem and IsobaricAnalyzer are available in Supplementary Table 3. Data are available via ProteomeXchange with identifier IPX0001849000.

### Differential Expression Profiles of Comparison H7N9 Infected with Healthy Control

We filtered the raw dataset (Supplementary Table 4), and identified 273 features which were only found in H7N9 infected cases (Supplementary Table 5). Those 273 features which were only found in H7N9 infected cases were applied proteins interaction analysis by String. The results showed 222 nodes with 798 edges, 7.2 average node degree and 0.426 Avg.Local clustering coefficient with protein-protein interaction (PPI) enrichment with *p*-value less than 1.0e-16 (Supplementary Table 6). 206 of 273 proteins were annotated to cell part (GO:0044464) from GO Component analysis, and 94 of 273 proteins were from extracellular region (GO:0005576). We then performed Venn diagram analysis on above two proteins list, and found 85 common proteins that can function on both cellular position of cell part (GO:0044464) and extracellular region (GO:0005576). The most of 94 proteins were annotated to immune related pathways by the Reactome database. The top 3 pathways were immune system (HSA-168256, 43 (45.7%)), innate immune system (HSA-168249, 37 (39.4%)) and neutrophil degranulation (HSA-6798695, 31 (32.98%)). The fourth and fifth pathways were metabolism of proteins (HSA-392499, 26 (27.7%)) and post-translational protein modification (HSA-597592, 20 (21.3%)), respectively. However, those 94 proteins could not be confirmed the source, may be generated from cell death or normally secreted from stressed cells. The remaining 7 singleton proteins were TNFSF12-TNFSF13, AC073610.3 (Novel Protein), C1RL, FBLN2, OLFML3, SBSN and C4orf54. They may play an important role in avian influenza infection, for instance, protein AC073610.3 acts as GTP binding and obsolete signal transducer activity, the most related GGA2 protein associated with avian influenza antibody titers[18]. There were still 51 proteins that were not annotated by the String system. Since our study focused on the extracellular biomarkers, we did not do further analysis on those proteins which were only found with H7N9 infected patients.

The dataset would be further filtered according to the proportion of the missing value (cut-off: 90%) and got the final qualified dataset for statistical analysis (Supplementary Table 7). We got 168 qualified features, and 140 (82.8%) proteins were annotated to the extracellular region (GO:0005576) from GO Component analysis. The qualified dataset was normalized and then applied one-way ANOVA with *post-hoc* tests, we identified 50 features and 4 clinical characters (LRT, URT, PF and TWC) with statistically significant difference (FDR < 0.05; Supplementary Table 8), and HPX (Hemopexin, P02790) was significantly different from each other (FDR = 0.0014, Earlier >> Control, Later >> Control and Earlier >> Later). We performed PPI analysis on those 50 statistically significantly different proteins with all avian influenza A virus proteins, interaction analysis shown 50 nodes, 231 edges, 9.24 average node degree and 0.513 Avg.Local clustering coefficient with PPI’s enrichment *p*-value less than 1.0e-16 (Figure 3), found that M1 and PB2 tightly linked with host’s HSPA8 (P11142, *p* = 0.0042) which plays a core role in the protein quality control system, ensuring the correct folding of proteins, and downregulated in H7N9 infected cases.

**Figure 3.**
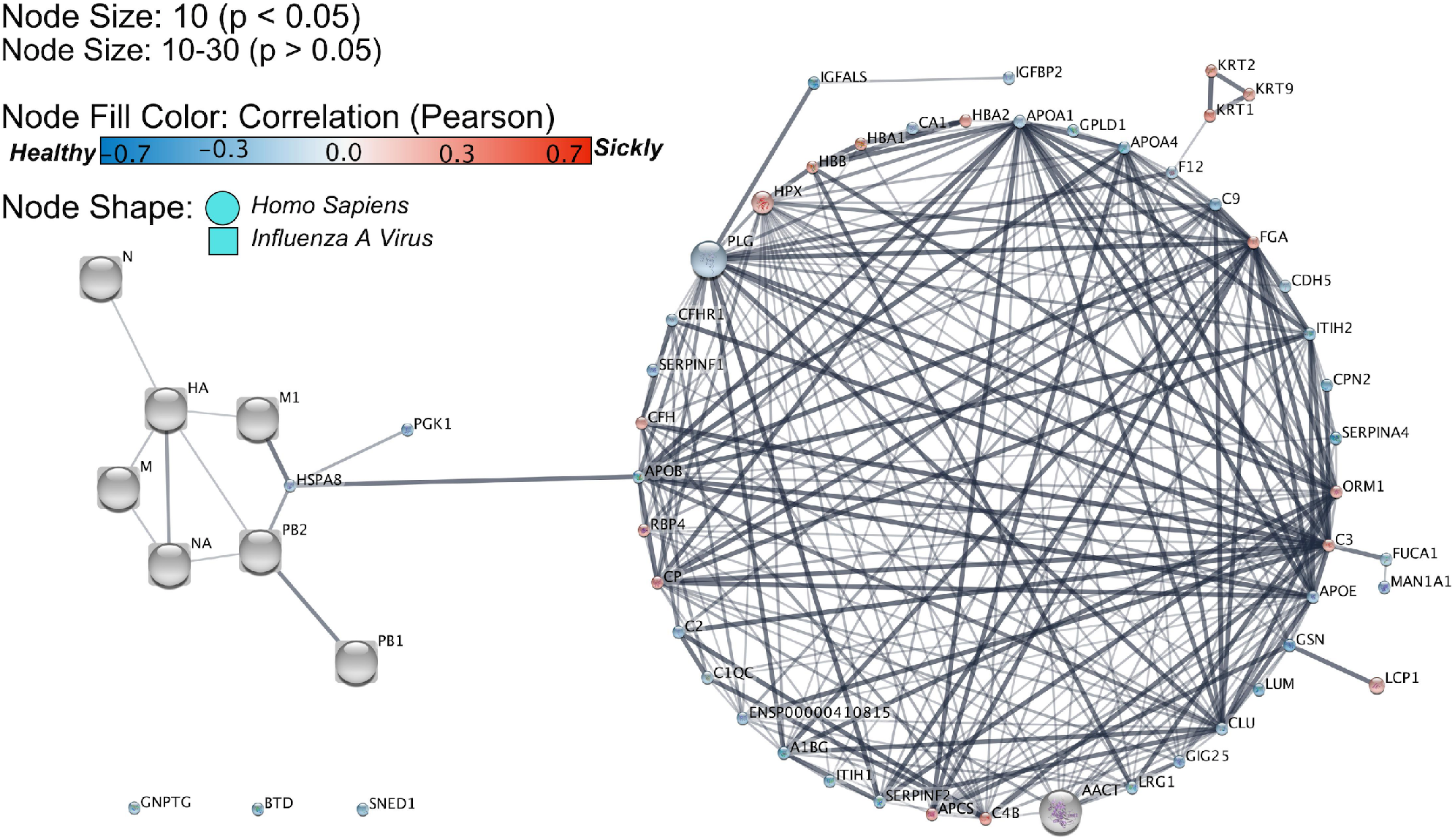
An interaction network between influenza A and extracellular host protein which were significantly different compared to healthy control. We downloaded influenza A proteins from public database using String, and we plotted by using Cytoscape (http://cytoscape.org/). Ellipse and round rectangle nodes indicate human proteins and influenza viral proteins, respectively. Node size is 10, means the protein is significantly correlated with disease status, and the node size is between 10 to 30 which means p-value is greater than 0.05, and positively correlated with the node size, node color is red/blue, means the protein positive/negative correlated with H7N9 infectious disease.

The biological function enrichment analysis on those 50 statistically significantly different proteins showed 44 proteins (88.0%) were annotated to extracellular region (GO:0005576) from cellular component analysis, 22 proteins (44.0%) were annotated to immune system (HSA-168256) from Reactome Pathways, 11 proteins (22%) were hit the complement and coagulation cascades pathway (hsa04610) from KEGG pathways annotation, and 9 proteins (18%) were enriched into endopeptidase inhibitor activity (GO:0004866) from GO Function, loss or reduction of endopeptidase inhibitor activity has been reported in vivo in influenza-infected ferrets[19]. The whole functional enrichment output was tabulated in supplementary table 9.

Correlation analysis can be carried out either against features or against a given pattern. The same qualified dataset (Supplementary Table 7) was applied correlation and pattern analysis with the Pearson method (FDR < 0.05, | *r* | > 0.5). The correlation of features with the values of different clinical characters, including PF ratio (PaO_2_/FiO_2_), procalcitonin (PCT), viral loads in the lower (LRT) and upper respiratory tract (URT), leucocyte (TWC), c-reactive protein (CRP), oxygen saturation (SPO_2_) and age, were performed by Pearson correlation test (Supplementary Table 10, 11 and Table 2). The total leucocyte (TWC) significantly negatively correlated with APOF (Q13790, *r* = − 0.54), APOA1 (P02647, *r* = −0.54) and C1QC (P02747, *r* = −0.53), and positively correlated with KRT1 (P04264, *r* = 0.50). Oxygen saturation (SPO_2_) level has been well confirmed that is associated with lung pathology in influenza infection. SPO_2_ level negatively correlated with C4BPA (P04003, *r* = − 0.53), F13A1 (P00488, *r* = −0.51), A2M (P01023, *r* = −0.55), PCSK9 (Q8NBP7, *r* = −0.51), PLTP (P55058, *r* = −0.52), APOF (Q13790, *r* = − 0.56), APOA1 (P02647, *r* = −0.58) and C1QC (P02747, *r* = −0.59), positively correlated with VTN (P04004, *r* = 0.52), APCS (P02743, *r* = 0.52) and LTF (P02788, *r* = 0.50). To investigate those features correlated with oxygen saturation level, we performed biology functional analysis on those proteins, and found they were highly linked together, and most of them were related with regulation of acute inflammatory response (GO:00002673), regulation of response to external stimulus (GO:0050727) and regulation of defence response (GO:0031347). The PF ratio significantly correlated with CKM (P06732, *r* = 0.50), MBL2 (P11226, *r* = 0.50), CA1 (P00915, *r* = 0.50), CDH5 (P33151, *r* = 0.50), APOE (P02649, *r* = 0.52), CLU (P10909, *r* = 0.54), ITIH2 (P19823, *r* = 0.52) and PLG (P00747, *r* = 0.51). We also performed biology functional analysis on those features and found that those proteins enriched into regulation of response to external stimulus (GO:0032101) and positive regulation of amyloid fibril formation (GO:1905908). Accumulation of beta-amyloid is a clinical sign of Alzheimer’s disorder, and Mitchell R. reported beta-amyloid inhibits replication of seasonal and pandemic flu[20].

**Table 2:**
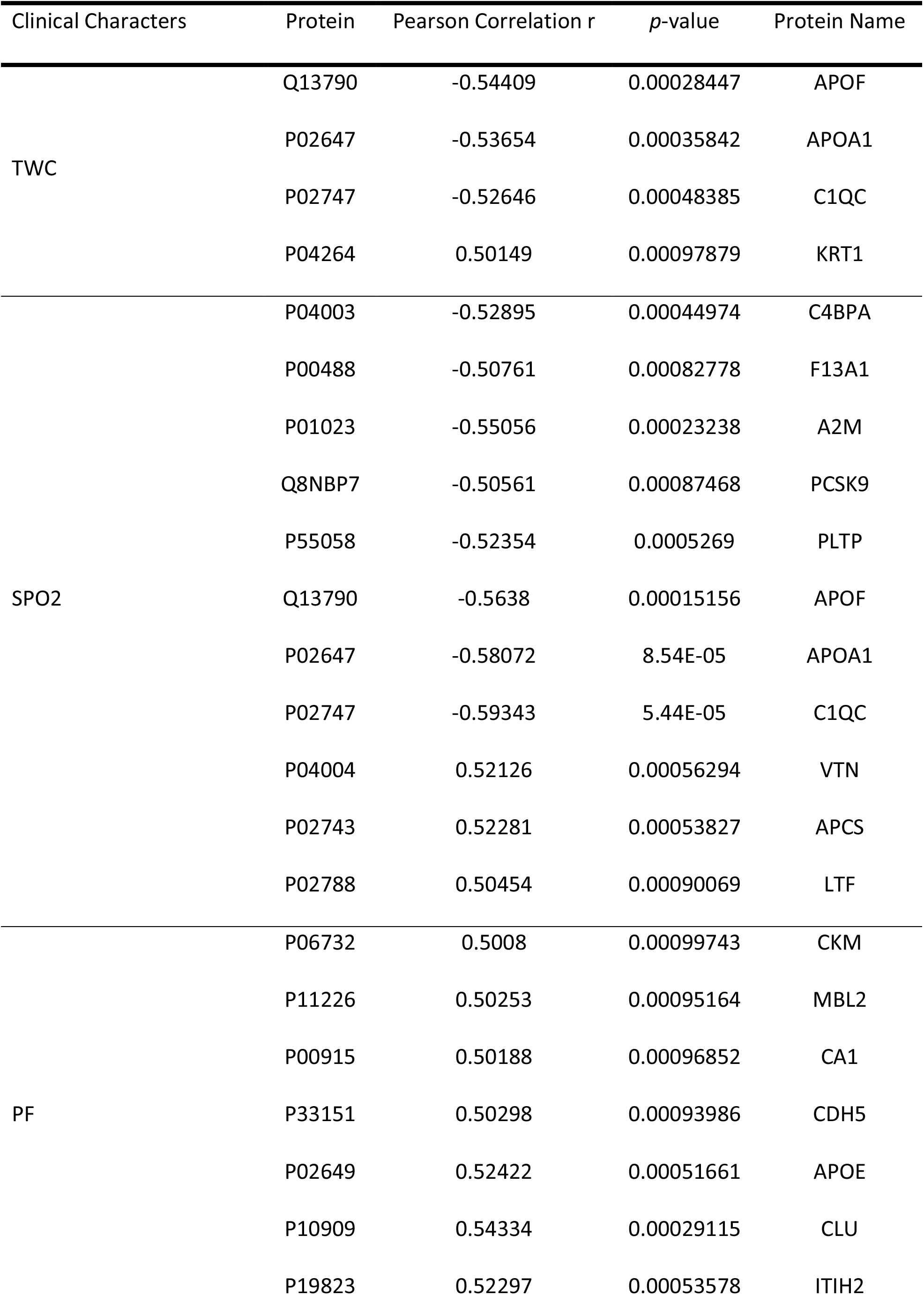
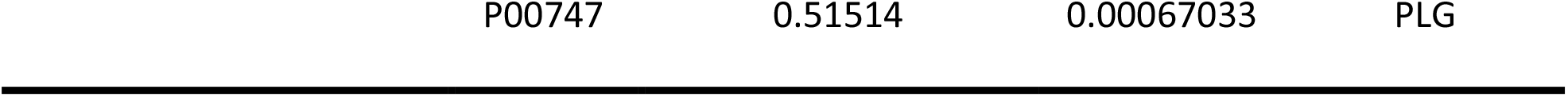
Sensitivity of proteins to clinical characters

Aim to identify whether the proteins or clinical characters correlated with H7N9 infection. Pattern analysis was performed on the same dataset (Supplementary Table 7), and termed “1-2-3” as the pattern used to search for interesting features that linearly increase with three groups: (Control-Earlier-Later). We identified 112 proteins and 2 clinical characters (URT and PaO_2_/FiO_2_ ratio) correlated with healthy control status (*r* < 0), and the rest proteins and clinical characters linked to H7N9 infectious status (*r* > 0) (Supplementary Table 12). Sixteen proteins and 1 clinical character (TWC) significantly upregulated in H7N9 infection (*r* > 0.39, FDR < 0.05), and PPI’s enrichment analysis showed two separately bins, one bin includes KRT family, KRT1, KRT2 and KRT9, the rest proteins clustered to the 2^nd^ bin. Thirteen of 16 proteins annotated to metabolic process (GO:0008152) by Biological Process database, and 5 of 16 proteins enriched into regulation of protein activation cascade (GO:2000257). FGA (P02671, Fibrinogen alpha chain), significantly positively correlated with H7N9 infectious status (*r* = 0.53, *p* = 0.0068). Fibrinogen was upregulated in a non-specific way in acute inflammatory conditions (infections, rheumatoid arthritis and other inflammations, cancers, etc.). There were 33 proteins significantly downregulated in H7N9 infection status (*r* < −0.39, FDR < 0.05), and biological function analysis showed those proteins enriched into regulation of proteolysis (GO:0030162), protein processing (GO:0070613) and inflammatory response (GO:0050727).

## Discussion

Clinical laboratory examinations are important for assessing the condition of H7N9 infected patients, however, it is not enough to reveal the etiologic factors that contribute to clinical phenotypes. We tested several clinical factors, including the viral load on upper/lower respiratory tract, the total white cell counts, c-reactive protein, procalcitonin, oxygen saturation (SPO_2_) and PaO_2_/FiO_2_ ratio, which have been correlated with influenza virus infection[3,21]. Influenza virus infection may cause host cells death via several different ways, including apoptosis[22], necroptosis[23] and pyroptosis[24]. H5N1 causes mammalian airway epithelial cells death via apoptosis[25]. A study of H7N9 infected CD14^+^ monocytes showed activation of caspases−8, −9 and −3 at 6h post infection, indicating that apoptosis and necroptosis pathways were simultaneously activated[26]. Our data showed that 42.1% of features (273) were only detected in H7N9 infected cases, and 75.5% (206/273) were annotated into cell part (GO:0044464) from GO Component analysis. Among those cell part (GO:0044464) proteins, still have 84 proteins were also expressed in extracellularly (GO:0005576), and biological function enrichment results showed most of them were annotated into immune response related pathways by Reactome database. These proteins might be generated by cell leakage/death due to H7N9 infection. High levels of cell component leakage can aggravate inflammation, and exacerbated the host immune responses, causing lung pathogenesis leading to acute respiratory distress syndrome (ARDS) with lower PaO_2_/FiO_2_ ratio, which correlated with disease severity level (Figure 4). Due to the limitations of study design, we could not confirm those proteins source, Therefore, they may have been generated from cell death or normally secreted/leaked from stressed cells.

**Figure 4.**
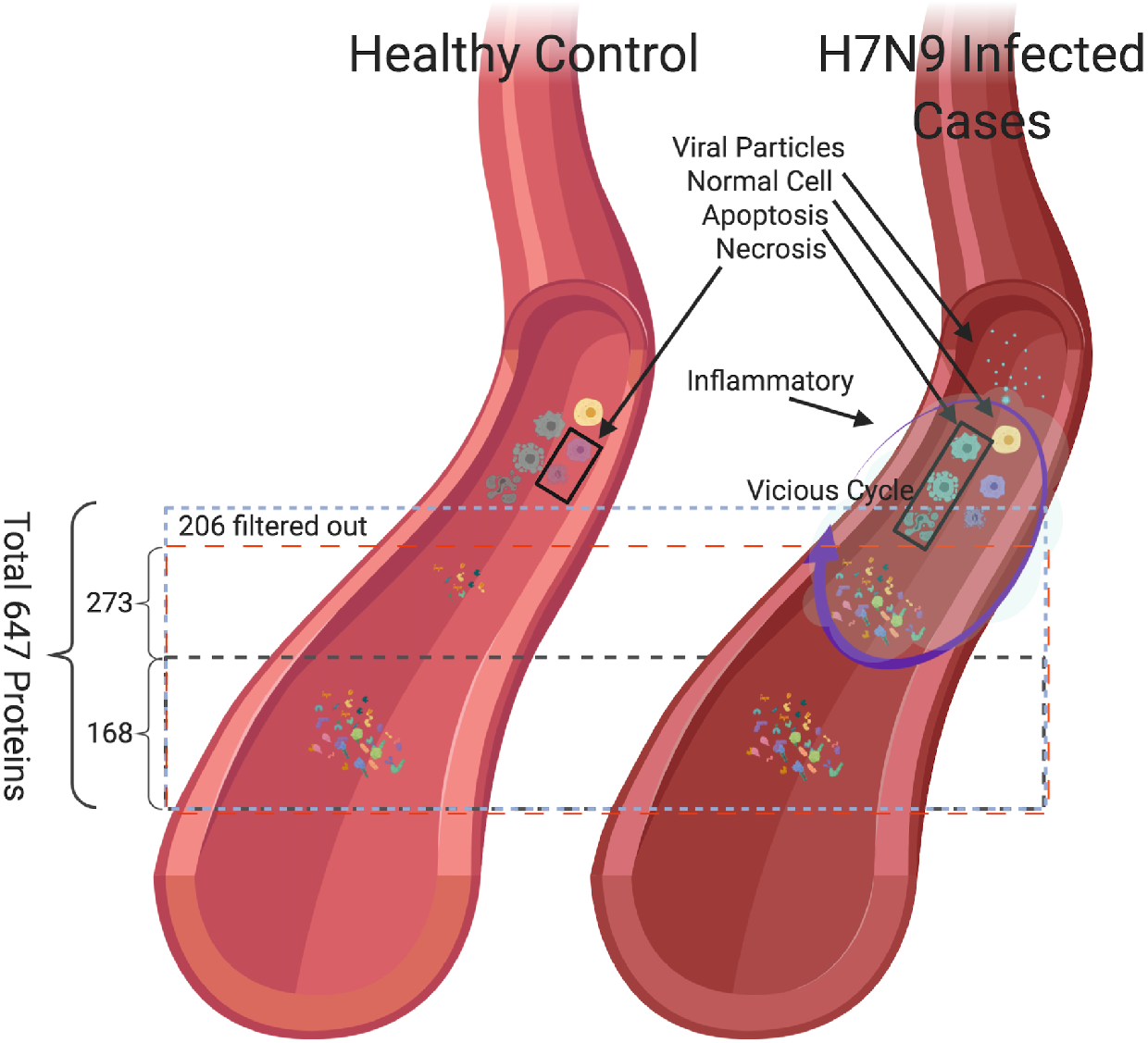
Summary of H7N9 infected and healthy control sera proteomics profiling. In total, we identified 647 proteins, and we dropped 206 proteins with more than 50% missing values in order to get more reliable results, and also identified 273 proteins were only found in H7N9 infected cases which might generate from cell leakage/death (apoptosis and/or necrosis), high levels of cell component leakage can aggravate inflammation, and exacerbated host immune responses caused lung pathogenesis.

The Secretory proteins and extracellular vesicles may act via cell-cell communication, delivery hormones, enzymes, and antimicrobial peptides. Extracellular fluid proteomic profiling might help understanding the host molecular response to influenza and identifying the novel prophylactic and therapeutic host biomarkers. In this study, we identified three heat shock proteins, two from the HSP70 family (HSPA8, HSPA5) and one from the HSP90 family (HSP90B1). The level of HSPA8 in serum may vary in certain pathological states. HSPA8 is widely expressed in the plasma membrane, extracellularly, in the nucleus and cytosol and the secreted/extracellular HSPA8 has important metabolic roles[27], acting as a trigger to boost pro-inflammatory cytokines expression, including TNF-alpha, IL-1beta and IL6[28]. HSPA8 may also regulate the hepatitis C virus replication (via both positive and negative ways)[29]. Kobayashi’s group firstly confirmed that HSPA8 interacts with Influenza virus matrix protein 1 (M1)’s C terminal domain, which plays a key role in the virion replication, assembly and maturation. The same research group down-regulated HSPA8 with siRNA treatment results in reduction of viral load which inhibited the viral M1 and vRNP protein’s nuclear exportation. They also demonstrated that HSP90 associated with the M1 and PA protein, concluding that HSPA8 and HSP90 may play important roles in various steps of the viral replication, assembly and maturation[30]. Therefore, Regulating the level of HSPA8 might be a therapeutic host biomarker for treatment of influenza infectious disease. Judith Frydman group reported that HSP70 family proteins have been shown to be involved into ZIKV virus replication, and HSP70s inhibitors are significantly reduced ZIKV replication in human and mosquito cells without appreciable toxicity, suggesting that HSP70s inhibitor can be used to treat the emergence of drug-resistant virus[31]. However, our result showed that HSPA8 was significantly downregulated in H7N9 infected cases (*r* = −0.39, FDR = 0.045), and HSPA8 correlated with APOA1 (P02647, *r* = 0.35), APOE (P02649, *r* = 0.26) and CLU (P10909, *r* = 0.11), which were annotated into amyloid-beta binding (GO:0001540, FDR = 0.0043). Pattern analysis results showed that APOA1, APOE and CLU were negatively correlated with H7N9 infection status with *r* = −0.38, FDR = 0.051; *r* = −0.46, FDR = 0.017 and *r* = − 0.41, FDR = 0.036, respectively. APOE is the major genetic risk factor for Alzheimer’s disease (AD)[32], and regulation of beta-Amyloid (βA) metabolism, aggregation and deposition. Upregulation of HSP70/HSP90 has also been shown to slow down Alzheimer disease development related to βA aggregation[33]. βA has been shown to inhibit growth of bacteria and fungi, and the replication of seasonal and pandemic strains of H3N2 and H1N1 influenza A virus in vitro[20]. Our study suggests that HSPA8, through its role in activating the amyloid-beta binding network, may be important in limiting the pathogenesis of H7N9 infection, although this remains to be proven in future study (Figure 5). A new compound, SW02, which can specifically upregulated HSP70s has already been identified[34,35], which may be an entirely novel way to therapeutic intervention of H7N9 infection.

**Figure 5.**
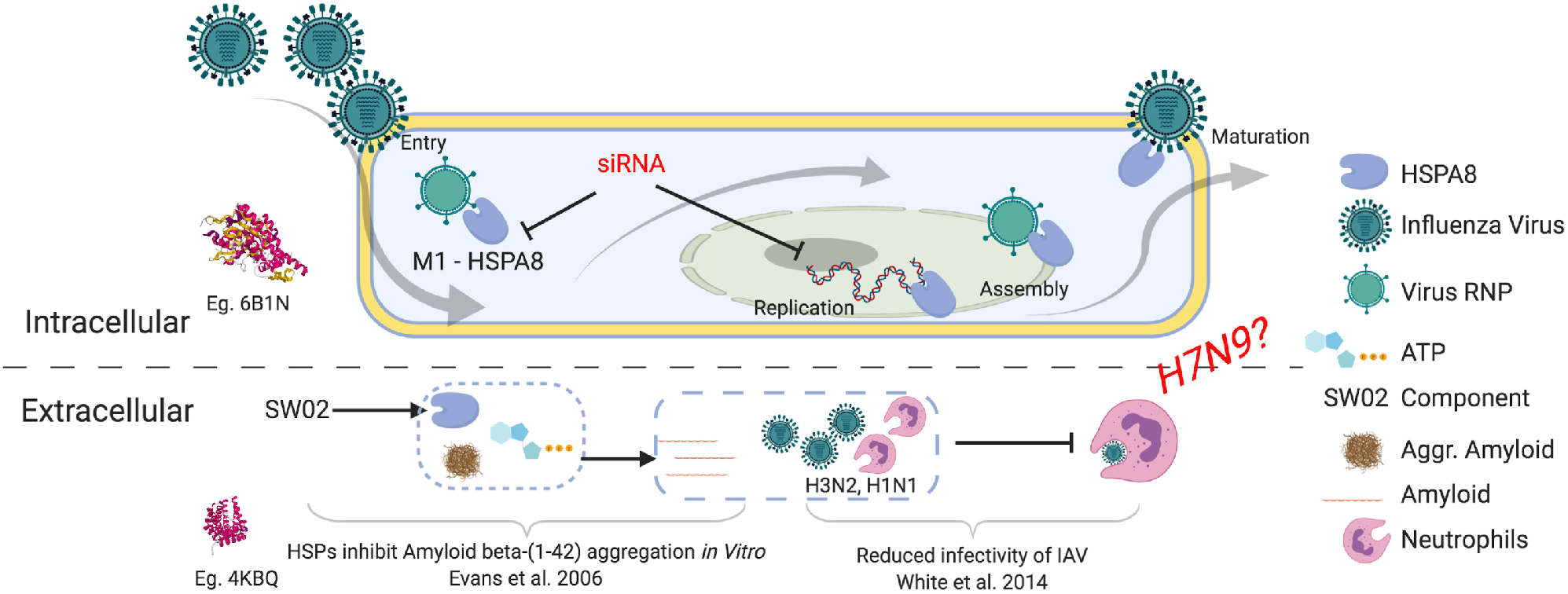
Summary of heat shock protein HSPA8 regulate the influenza A virus replication via both positive and negative way. Intracellular: Down-regulated HSPA8 with siRNA treatment results in reduction of influenza virus load which inhibited the viral M1 and vRNP proteins nuclear exportation. Extracellular: Upregulated HSPA8 with SW02 (HSPA8 activator) may active amyloid-beta binding network, then the activated network regulates the beta-amyloid protein which has been reported that suppress the H3N2 and H1N1 virion replication, assembly and maturation. Our study suggests that it may also work for suppression H7N9.

In summary, population-based extracellular fluid proteomic profiling was performed on 15 H7N9 infected cases compared to 10 healthy controls. We identified features that were distinct only in H7N9 patients, and most of those features may be from cell leakage/death. We also identified novel candidates that can serve as potential prophylactic and therapeutic host biomarkers. Due to limited new H7N9 cases (as of October 2017 to now, there were only 4 additional cases[35]), we did not verify the findings in the same sample sets or apply the verification study on the independent sample sets. This has limited our study.

## Acknowledgments

We would like to thank all the clinical staff and research nurses who contributed to this study. We would like to thank Dr. Sook-San Wong and Dr. Mark Zanin for their suggestions on the article.

